# An automated framework for efficiently designing deep convolutional neural networks in genomics

**DOI:** 10.1101/2020.08.18.251561

**Authors:** Zijun Zhang, Christopher Y. Park, Chandra L. Theesfeld, Olga G. Troyanskaya

## Abstract

Convolutional neural networks (CNNs) have become a standard for analysis of biological sequences. Tuning of network architectures is essential for CNN’s performance, yet it requires substantial knowledge of machine learning and commitment of time and effort. This process thus imposes a major barrier to broad and effective application of modern deep learning in genomics. Here, we present AMBER, a fully automated framework to efficiently design and apply CNNs for genomic sequences. AMBER designs optimal models for user-specified biological questions through the state-of-the-art Neural Architecture Search (NAS). We applied AMBER to the task of modelling genomic regulatory features and demonstrated that the predictions of the AMBER-designed model are significantly more accurate than the equivalent baseline non-NAS models and match or even exceed published expert-designed models. Interpretation of AMBER architecture search revealed its design principles of utilizing the full space of computational operations for accurately modelling genomic sequences. Furthermore, we illustrated the use of AMBER to accurately discover functional genomic variants in allele-specific binding and disease heritability enrichment. AMBER provides an efficient automated method for designing accurate deep learning models in genomics.

## Main

Artificial neural networks, or deep learning, have become a state-of-the-art approach to solve diverse problems in biology^1,2^. Convolutional Neural Networks (CNNs) are especially well-suited for identifying high-level features in raw input data with strong spatial structures^3^ and as such are powerful at modelling raw genomic sequences and extracting functional information from billions of base-pairs in the genome^1^. CNN-based approaches address the computational challenges of predicting the chromatin state and RNA-binding proteins binding state from sequence^4–6^, identifying RNA splice sites^7^, predicting gene expression^8^, and prioritizing disease relevance of variants^9^, and many more^1^. Overall, CNNs have become the de-facto standard for analysis of genomes - a fundamental problem in both basic understanding of biology and for enabling personalized and precision medicine approaches.

The successful applications of CNNs have been largely attributed to their corresponding architectures. Indeed, for CNN applications in genomics and biomedicine, numerous efforts have been devoted to the development of architectures, such as in DeepSEA^4^, Basenji^10^ and SpliceAI^7^. This is similar to the extensive efforts in architecture designs for tackling computer vision problems, for example VGG^11^, Inception^12^, and ResNet^13^. Each of these architectures is motivated and inspired by deep understanding of machine learning and domain knowledge; and requires substantial effort and time commitment by experts to design and implement by extensive trial-and-error processes.

Here, we present Automated Modelling for Biological Evidence-based Research (AMBER), an automatic framework for efficiently designing convolutional neural networks in genomics. To our knowledge, AMBER is the first automated approach specifically designed for modelling genomic sequences. It leverages the groundbreaking idea of Automated Machine Learning (or AutoML), and the related family of algorithms for Neural Architecture Search (NAS) previously developed in the context of computer vision^14,15^. For a given fixed set of training data, AMBER designs an optimal architecture by NAS in a pre-defined model space. We show that the AMBER-designed models significantly outperformed equivalent non-NAS models, matching or even exceeding published expert-designed models. Finally, we use two well-established benchmarks to demonstrate that the AMBER-designed optimal architectures provided significant advantages in prioritizing functional genomic variants in allele-specific binding and heritability enrichment in Genome-Wide Association Studies (GWAS). We also illustrate the use of AMBER-designed models to discover disease-relevant variants. Thus, AMBER creates accurate and informative deep-learning models that can support functional genomics discoveries by biologists with and without machine learning expertise. AMBER is publicly available at https://github.com/zj-zhang/AMBER.

## Overview of methods and workflow

The AMBER framework fully automates the process of training and applying deep learning to genomics, including automatic design of neural network architecture from the training data and downstream functional analyses with the AMBER-designed model (Figure 1). Unlike existing approaches that focus on making deep learning more accessible using established model architectures^16,17^, AMBER automatically designs an optimal architecture for each user-specified problem.

**Figure 1.**
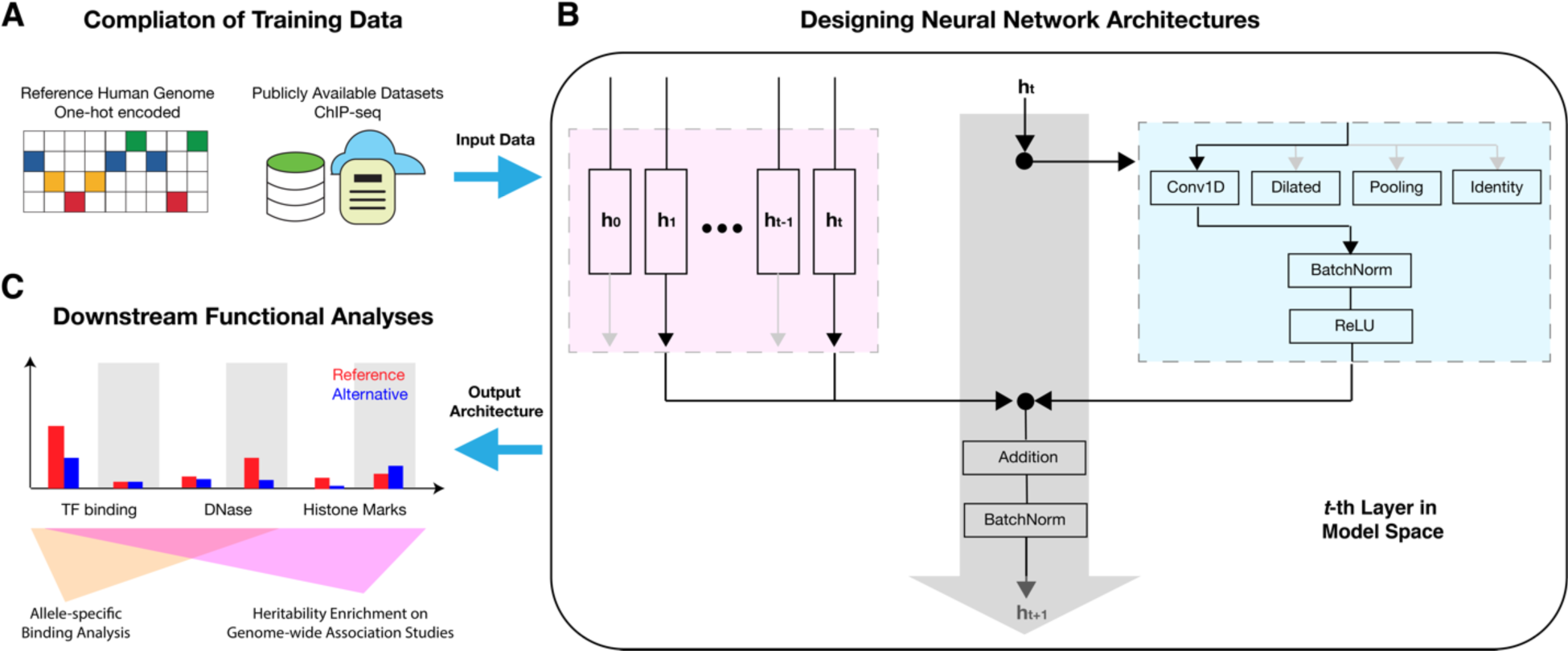
Method and workflow overview of AMBER. **A)**AMBER uses a compendium of training data to design deep learning models in functional genomics. In this application, we applied AMBER to the task of predicting transcriptional regulation on DNA sequences. The features are one-hot encoded reference human genome, and the labels are functional annotations derived from a large set of ChIP-seq data. **B)** AMBER designs network architecture by searching for optimal combinations of computational operations (blue box) and residual connections (red box) for each layer, to construct a child model that maps training features to training labels. **C)** Taking the optimal architecture as output, AMBER performs downstream functional analyses. For the transcriptional regulation model, we analyzed the functional variant prioritization by AMBER-designed models to predict allele-specific binding and heritability enrichment in GWAS.

In general, to investigate a biological question with AMBER, a biologist would compile a compendium of functional genomics data such as profiles of transcription factor binding or histone marks along the genome. AMBER uses such sets of compiled training features and labels as input to automatically design deep learning models for the biological question or task of interest (Figure 1A). Here, we use AMBER to model transcriptional regulatory activities. For this task, the training features are one-hot encoded matrices that each represent 1000-bp DNA sequences from the reference human genome, and the training labels are binary outcomes derived from a compendium of 919 distinct transcriptional regulatory features. These regulatory features include four main functional categories in diverse tissues and cell lines: transcription factors (TF), polymerases (Pol), histone modifications (Histone), and DNA accessibility (DNase). The task aims to predict whether one or more of the 919 transcriptional regulatory features are active for any 1000-bp human DNA sequences. In total, the training dataset spans more than 500 million base-pairs of the human genome, with 4400000, 8000, and 455024 samples for training, validation and testing, respectively. Conditioned on this dataset, the target model for AMBER to design is a convolutional neural network with multi-tasking consisting of 919 individual tasks.

To more formally define the neural architecture search problem, the target convolutional neural network architecture can be divided into two interconnected components: the computational operations used in each layer (blue box, Figure 1B), and the residual connections from previous layers (red box, Figure 1B). Residual connections have been demonstrated to enable the training of much deeper neural networks with superior performances^13^, while greatly expanding the model search space (7.4 × 10^19^ times more viable architectures in our model space; see **Methods**). Thus, it’s essential that residual connection search is considered when AMBER searches for architectures, and the search needs to be efficient. AMBER searches for both of the two components jointly using the Efficient Neural Architecture Search (ENAS) controller model^15^. The controller model is parameterized as a Recurrent Neural Network (or RNN; for details, see **Methods**). Briefly, for each layer in the model search space, the probability of selecting a computational operation is computed by a multivariate classification dependent on the current RNN hidden state; and the probability of selecting the residual connections from a previous layer is a function of the RNN hidden states of the current layer as well as the previous layer of interest. The RNN hidden states were subsequently updated by the operations or residual connections sampled from the output probabilities. To train the controller RNN, we employed reinforcement learning to maximize a reward of AUROC on the validation dataset.

The output of AMBER is an optimized architecture that performs better than architectures uniformly sampled from the same model search space (**Methods**). Furthermore, we show that AMBER-designed models provide significant advantages over baseline models in multiple practical scenarios, including allele-specific binding and heritability enrichment in GWAS. In the following sections, we describe each part of the AMBER pipeline as well as the downstream analyses in detail.

## AMBER designs accurate and efficient models

In our example AMBER application, we defined the model search space of 12 layers, each layer with 7 commonly used computational operations. We chose to use a 12-layer model space because this was the maximum hardware memory limit for a single Nvidia-V100 GPU, and shallower models can be attained by an identity operator that in effect removed one layer. In total, this model space hosts 5.1 × 10^30^ distinct model architectures (**Methods**).

We benchmarked the computational efficiency of AMBER by comparing the GPU time used by the AMBER search phase to other architecture search algorithms (Table 1). The time of AMBER search phase is orders of magnitude more efficient than RL-NAS^14^ and AmoebaNet^18^ and comparable to DARTS^19^ and ENAS^15^.

**Table 1.** Runtime comparison in GPU hours

| Method | Time (in GPU hours) |
| --- | --- |
| AMBER | 72 |
| AmoebaNet | 75600 |
| DARTS | 96 |
| ENAS | 10.8 |
| RL-NAS | 537600 |

To robustly evaluate the accuracy of AMBER-designed models, we performed six independent runs of AMBER architecture search, generating six “searched models”. We compared these searched models with uniformly sampled residual network architectures from the same model space (“sampled models”). Given the architectures, the final training step for AMBER architectures and the sampled residual network architectures were identical, with all models trained to convergence (**Methods**).

The average testing AUROC and AUPR for each functional category of 919 regulatory feature prediction tasks (i.e. TF, Pol, DNase and Histone) were compared for the six searched and six sampled model architectures. AMBER-designed architectures significantly outperformed the sampled architectures for all categories (Figure 2A). The prediction accuracies of different models were more alike within a given functional category than across different categories, indicating that the inherent characteristics of the training data play an essential role in the model’s prediction performance, regardless of its model architecture. This is expected, because the training data determined the upper bound of model performance^20^, while the searched architectures better approximated this bound. Of course, with unlimited time and resources to enable complete sampling, the optimal architecture is theoretically reachable by sampling as well; however, the time and resource consumption will be tremendous in a model space of 5.1 × 10^30^ potential architectures. The AMBER architecture search by far speeds up this process and yields model architectures in a narrow high-performance region. Detailed performances for each model can be found in **Supplementary Table 1**. Hence, AMBER robustly designs high-performance convolutional neural network architectures.

**Figure 2.**
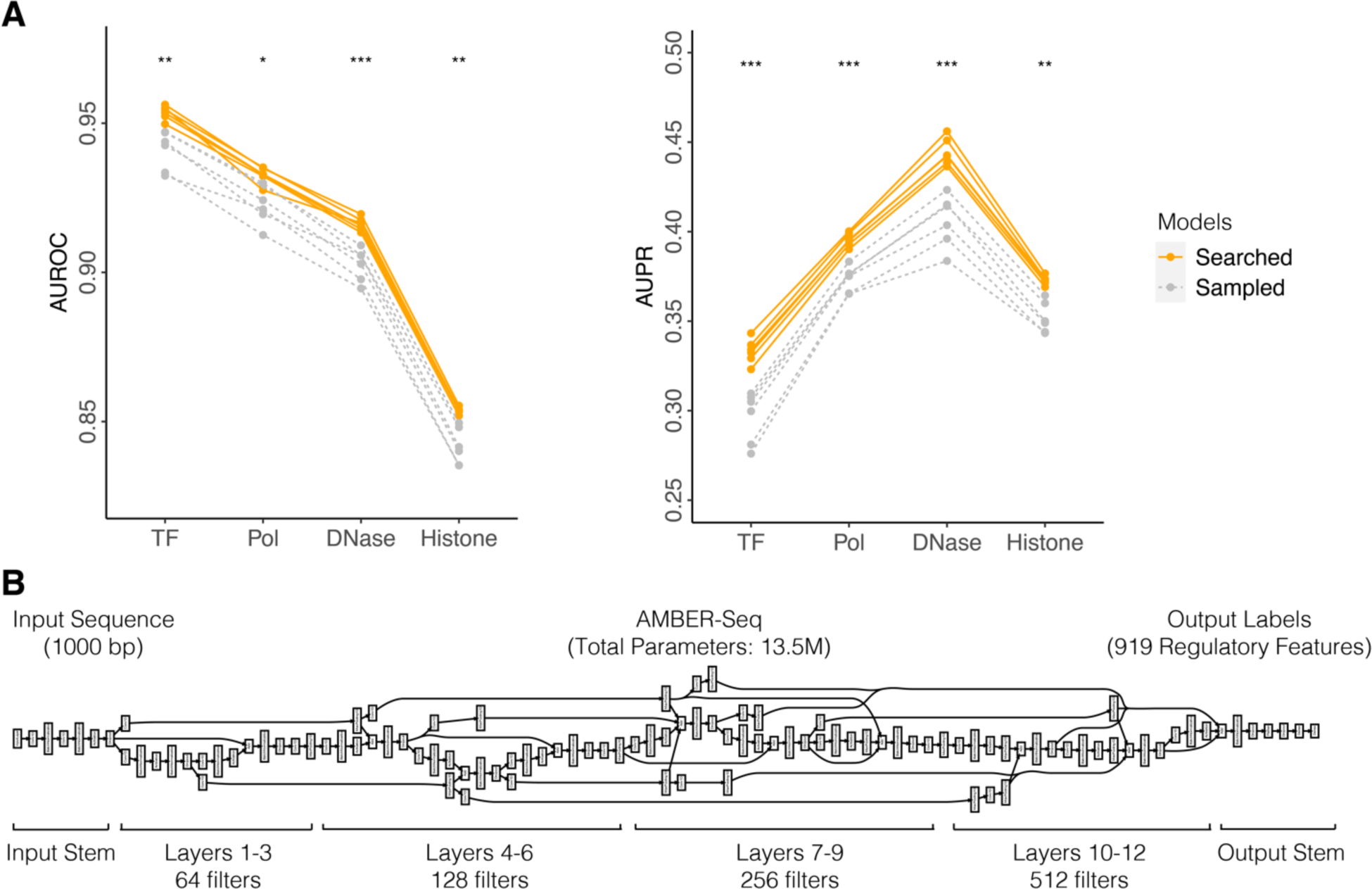
AMBER searched architectures outperform sampled architectures. **A)** The average testing AUROC and AUPR in each functional category were compared for twelve models with distinct architectures either generated by AMBER searched (orange) or uniformly sampled from model space (grey). Each model, represented by a line, was identically trained to convergence. **B)** An illustration of the optimal model architecture, AMBER-Seq, used for downstream analyses. AMBER-Seq is an AMBER-designed deep convolutional neural networks that outputs a multi-label binary classification for 919 transcriptional regulatory features using 1000-bp DNA sequences as inputs. Statistical significance (t-test) *: p<0.05, **: p<0.01, ***: p<0.001

Theoretically, the superior performance from searched model architectures could be achieved by higher relative model complexity. However, no significant differences were observed between the two groups of architectures (p-value=0.69, t-test). When we examined the total number of parameters in each child architecture (dot sizes, **Supplementary Figure 1**), the average number of parameters is 12.9 million for searched architectures and 13.3 million for sampled architectures, respectively. Furthermore, we did not observe correlations between the model complexities and their testing performances (spearman correlation=0.06, p-value=0.87). This indicates that the superior performance from searched model architectures is not explicitly linked to model complexities, and that AMBER-designed models are parameter-efficient.

For the rest of the analyses in this study, we used the AMBER-designed architecture with the best testing performance, referred to as AMBER-Seq (Figure 2B); and compared it to the sampled architecture with the best testing performance, referred to as AMBER-Base (**Supplementary Figure 2**). Starting with the 1000-bp one-hot encoded input, we use the input stem of one convolutional layer to expand the 4-channel DNA sequence into 64 channels. The input stem is identical for all child networks. Similarly, the output stem flattens the convolutional feature maps, followed by a dense layer of 925 hidden units to predict the 919 regulatory outputs. The middle 12 layers are variable and grouped into four blocks, each with 3 layers. The total number of parameters in AMBER-Seq is 13.5 million, which is substantially fewer than the original expert-based implementation (52.8 million) in ref.4 and a model of a similar task (22.8 million) in ref.^10^. With fewer total parameters, AMBER-Seq matched and even exceeded the previously expert-designed implementation in prediction accuracy (AUROC and AUPR; see **Supplementary Table 1**).

## Deciphering the logic of AMBER architecture search

Unbiased architecture search performed by AMBER provides insight into which computational operations and architectures are most suited for particular problems in genomics. This can diagnose whether the controller RNN model has learned meaningful representations and help design better model search spaces for future applications.

For this analysis, we analyzed the average probability of all computational operations in the last step of the AMBER-Seq controller training across the 12 layers (Figure 3A). The likelihood of using convolutions (vanilla and dilated convolution) was the highest in the bottom- to middle-layers; in particular, convolution with kernel size 8 was universally preferred, which is consistent with the choice in expert-based architectures^4^. Interestingly, in higher layers, the likelihood of max pooling starts to increase as the layers are closer to the output. In light of CNN’s hierarchical representation learning in computer vision^21^, we speculate this is because more high-level features with biological semantic meanings are constituted in the top layers of convolutions, after extensive usage of convolution operations in the bottom layers. Subsequently, by using max pooling as the computational operation in top layers, the model performs feature selections that regularizes model complexity and encourages the usage of high-level semantic features in predicting the final regulatory outcomes. We anticipate this AMBER architecture design pattern can be further generalized and transferable to other related tasks^22^.

**Figure 3.**
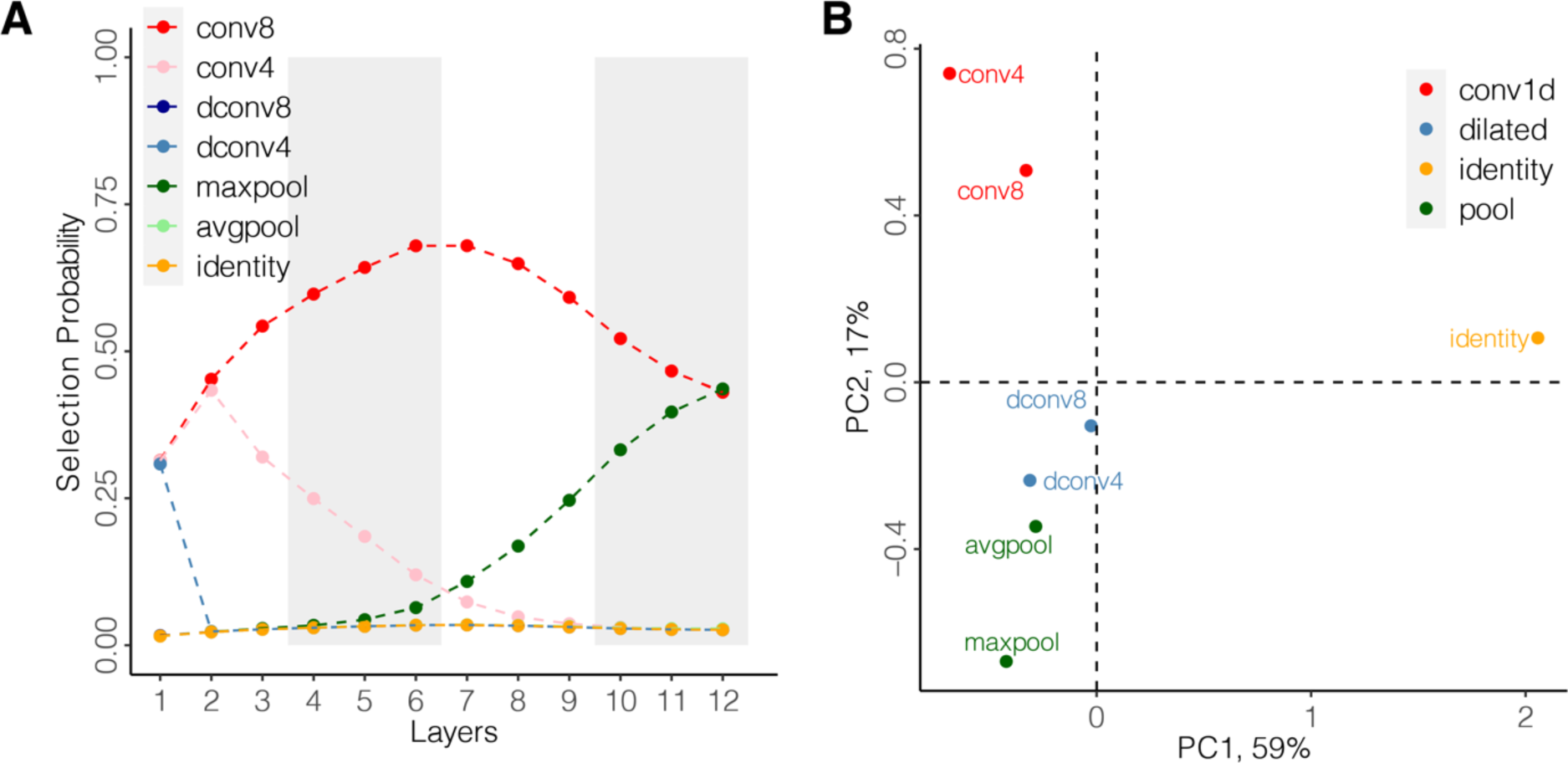
Illustration of AMBER architecture search logistics. **A)**Selection probabilities for distinct computational operations in each layer of the AMBER-Seq controller. For this architecture, convolutional operations were preferred in bottom to middle layers, while the likelihood of selecting max pooling increased in top layers. **B)** Principal component analysis of the embedding vectors for different computational operations. PC1 separated identity from computational operations; PC2 separated vanilla convolution, dilated convolution and pooling. Abbreviations: *conv8/4*: 1D convolution with kernel size 8/4; *dconv8/4*: dilated convolution with kernel size 8/4; *max/avgpool*: max/average pooling.

The controller’s ability to distinguish distinct and similar computational operations is critical for searching high-performance architectures. The differential selection likelihood of operations across layers is a function of previous RNN hidden states and the embedding vectors for each operation, which are learned during the AMBER search phase (**Methods**). We performed Principal Component Analysis (PCA) on the embedding vectors and analyzed how AMBER distinguishes operations (Figure 3B and **Supplementary Figure 3**). We found that the first principal component separates identity from all other computational operations, as the identity layer does not involve any computations. In the second principal component, convolution and pooling were separated with dilated convolution as an intermediate between vanilla convolution and pooling layers. Indeed, dilated convolution enlarges the receptive field similar to pooling layers, while also performs convolution computations^23^. The third principal component further separated computational operations by their corresponding operation types (**Supplementary Figure 3**). Overall, AMBER controller RNN can distinguish between similar but distinct operations in building the target architecture.

## Variant effect prediction on allele-specific binding

A key application of convolutional neural networks in genomics is to predict functional effects of genomic variants, i.e. a variant’s potential to disrupt an existing molecular mechanism or generate a new one. To investigate the variant effect prediction of different neural network architectures, we compared their ability to correctly predict allele-specific binding for 52,413 variants in 83 distinct transcription factors generated by ChIP-seq experiments^24^. These experiments measure the effect of specific alleles on binding of transcription factors, providing an independent evaluation set for our predictions. For comparison, in addition to AMBER-Seq and AMBER-Base, we included a set of commonly used models and motifs for scoring variant effects: expert-designed CNNs DeepSEA^4^ and DeepBind^6^, deltaSVM^25^, Jaspar^26^ and MEME^27^ (Figure 4A). For comparison across different models, variant scores were rank transformed to the range of [-1, 1] and AUROC was computed for each method’s ability to distinguish loss/gain-of-binding alleles versus neutral alleles (**Methods**). In general, machine learning methods (AMBER, DeepSEA, DeepBind, deltaSVM) predict variant effects significantly better than the motif-based methods (i.e. Jaspar and MEME). Importantly, AMBER-Seq’s performance matched or exceeded all other methods, including expert-designed architectures and the AMBER-Base model, demonstrating the power of automated architecture search (asteroid, Figure 4A).

**Figure 4.**
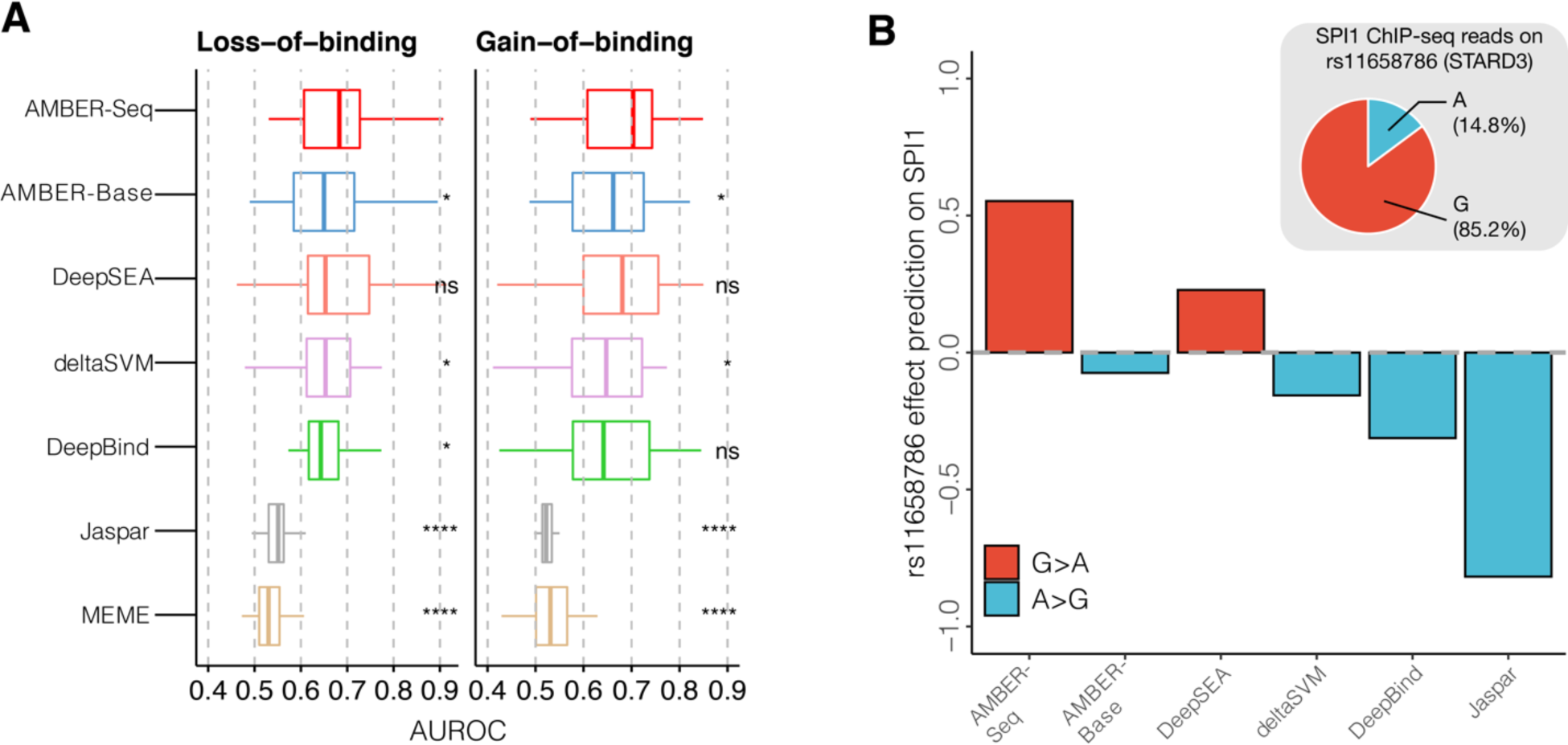
Benchmarking variant effect prediction with allele-specific binding. **A)**Performance of distinguishing loss- and gain-of-binding variants from different models and methods evaluated by AUROC. AMBER-Seq outperformed AMBER-Base on the compendium of allele-specific transcription factor binding sites, matching or even exceeding previous expert-designed machine learning methods. In each boxplot, center line marks the median while top and bottom lines mark the first and third quartiles. **B)** A biological case study of variant effect prediction of human genomic variant rs11658786. This variant was predicted to alter a SPI1 binding site in gene STARD3. Among different methods, only AMBER-Seq and DeepSEA predicted the loss-of-binding effect (G>A) of this variant. The A allele significantly reduces SPI1 binding, as illustrated by an independent ChIP-seq experiment (inset). Statistical significance of results of AMBER-Seq versus each of the other models (Wilcoxon test) ns: p>0.05, *: p<0.05, **: p<0.01, ***: p<0.001, ****: p<0.0001

As a biological case study, we focused on the effect of genomic variant rs11658786 on binding of the SPI1 transcription factor (Figure 4B). SPI1 (also known as PU.1) is a transcription activator with important functions in hematopoiesis^28^, leukemogenesis^29^, and adipogenesis^30,31^. AMBER-Seq predicted that the alternative allele at this position reduces SPl1 binding, a prediction supported by independent experimental data -167- in an independent ChIP-seq dataset, SPI1 predominantly binds to the G allele (85.2%) than the A allele (14.8%; Figure 4B, **inset**). Interestingly, all other models except DeepSEA predicted that the alternative allele enhances SPl1 binding, contradicting experimental results. Moreover, rs1165876 is an eQTL for its target gene, STARD3 (**Supplementary Figure 4A**), where the gene expression for the G genotype is the highest and the A genotype is the lowest. The eQTL effect for gene expression is consistent with the AMBER-Seq predicted effect of SPI1 binding and its transcription activation function. Finally, STARD3 is a gene that encodes a member of a subfamily of lipid trafficking proteins that is involved in cholesterol metabolism. By querying GWAS catalog^32^, we confirmed that rs11658786 is in strong LD with significant GWAS loci in high cholesterol, its interaction terms, as well as smoking status (**Supplementary Figure 4B**). Overall, this case study illustrates how variant effects accurately predicted by the automatically generated AMBER-Seq model can be useful for prioritizing functional variants of interest.

## Heritability enrichment analysis of genome-wide association studies

Finally, we assessed the utility of automatic architecture search by comparing AMBER-Seq with the uniformly sampled AMBER-Base model for explaining disease heritability in GWAS from UK Biobank^33^. Using AMBER-Seq and AMBER-Base models, variant annotations for each of the 919 transcriptional regulatory features of each model were generated, followed by stratified LD-score regression^34^ to evaluate their heritability enrichment for a given GWAS (**Methods**). We analyzed the GWAS summary statistics of disease phenotypes previously reported^35^ (**Methods**). The union of the significantly enriched variant annotations (FDR<0.05) from both models were used for downstream comparisons and were subsequently examined for overlapping between the AMBER-Seq and AMBER-Base models, or unique to either one of the models (**Supplementary Figure 5**). Of the six GWAS diseases we studied, five have significantly more enriched heritability in AMBER-Seq variant annotations (Figure 5A; **Methods**). On average, AMBER-Seq variant annotations were 1.81x more enriched in heritability compared to their counterparts in AMBER-Base across all diseases, indicating that AMBER-designed model produced more informative variant effect predictions for interpreting disease-associated genomic loci.

**Figure 5.**
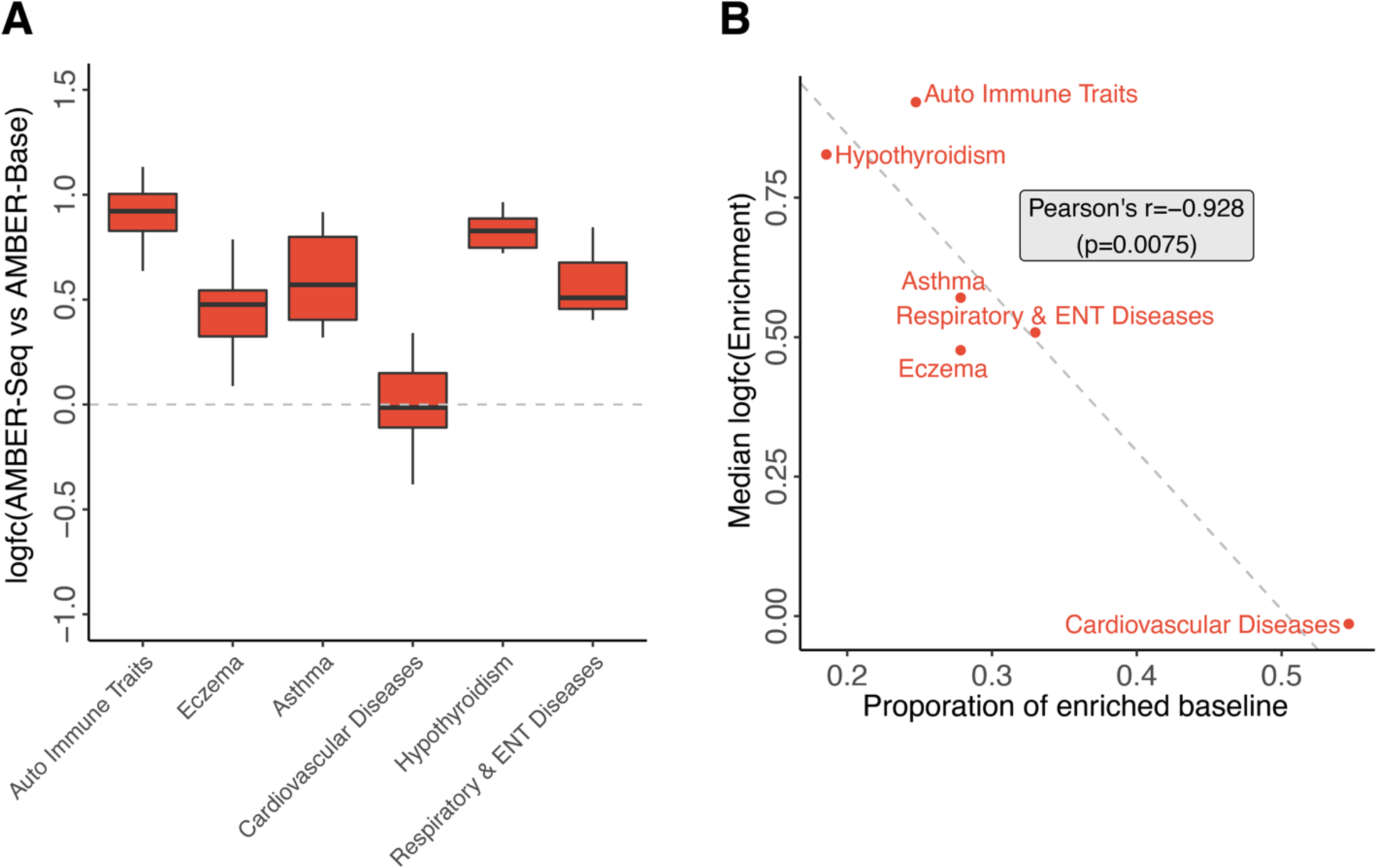
Benchmarking heritability enrichment in disease GWAS. **A)**Comparison of heritability enrichment of AMBER-Seq and AMBER-Base’s variant annotations for six disease GWAS. On average, AMBER-Seq annotations were 1.81x more enriched in disease heritability than the annotations of AMBER-Base. In each boxplot, center line marks the median while top and bottom lines mark the first and third quartiles. **B)** The median magnitude of enrichment fold-change between AMBER-Seq and AMBER-Base was negatively correlated with the proportion of enriched baseline annotations in various diseases, indicating that AMBER can deliver more informative variant annotations in diseases with poor baseline annotations.

Moreover, the variant annotations from AMBER-Seq were particularly useful where baseline annotations^34^ fail to explain heritability (Figure 5B). Baseline annotations are a collection of 97 functional annotations previously curated^34^ that cover major known regulatory patterns for human genome. Specifically, to quantify how well the baseline annotations alone explained heritability, we regressed baseline annotations for each GWAS phenotype and calculated the proportion of baseline annotations that were significantly enriched in heritability. We observed a significant negative correlation between the median log fold-change of heritability enrichment of annotations from AMBER-Seq over AMBER-Base, versus the proportion of baseline annotations that are significant (Figure 5B). This demonstrates that for disease where only a few baseline annotations were significantly enriched in heritability, AMBER-Seq provides the most improvement over AMBER-Base in variant annotation. Conversely, when AMBER-Seq and AMBER-Base heritability enrichment was comparable, the majority of the heritability was largely explained by baseline annotations. Therefore, the automated model design pipeline of AMBER is able to deliver more informative variant annotations in the cases where they are arguably most needed, i.e. for diseases that are poorly annotated by baseline annotations.

## Discussion

The past decade has witnessed a revolutionary transformation in genomics and exponential accumulation of high-throughput sequencing data. These data enable the study of diverse molecular mechanisms and biological systems through a quantitative lens. Deep learning models have been especially powerful in modeling biological sequences, transforming our ability to interpret genomes^4–6^. These methods generally employ convolutional neural networks to extract features from raw genomic sequences, but such an approach comes with a price: a convolutional layer has more hyperparameters than a regular fully connected layer, making the hyperparameter tuning a significantly harder problem. To date, the vast majority (if not all) of the deep learning models are manually tuned by computational biologists through trial-and-error, which is time consuming and imposes a substantial barrier for applications of such models by biomedical researchers. To address this challenge, we developed an automatic architecture search framework, AMBER, for efficiently designing optimal deep learning models in genomics. In this study, we have applied AMBER to predicting genomic regulatory features, including downstream analyses such as variant effect prediction and heritability enrichment in GWAS. We found that AMBER matched or exceeded performance of baseline models, including both expert-designed and uniformly sampled architectures, and is computationally efficient. We anticipate that AMBER will provide a useful tool for biomedical researchers, with and without machine learning expertise, to rapidly develop deep learning models for their specific biological questions.

An important additional application of AMBER is for upgrading existing models with advanced model architectures or updating models when additional data become available. Compared to the original implementation of DeepSEA in 2015, it is interesting to observe that all six runs of AMBER searched models performed better (**Supplementary Table 1**). This is especially relevant as new and powerful architectures are being developed continuously (e.g. residual connections^13^ that likely contribute to AMBER-Seq’s high performance), yet it is non-trivial to adapt models with the latest deep learning techniques, and such adoption is time- and effort-consuming. AMBER enables readily integrating such modern approaches into existing expert-designed models. With AMBER, researchers can easily build and apply modern deep learning techniques to find the optimal neural architecture, thereby accelerating the scientific discoveries in biology.

Finally, an important future direction for architecture search in biology is to jointly optimize the prediction accuracy as well as model interpretability. For example, elucidating the decision logic behind variant prediction can help identify molecular pathways that likely led to the predicted effects, shedding new light on molecular mechanisms of transcriptional regulation^36^. In general, an interpretable model is particularly desirable when practitioners need explicit evidence for decision making and/or for knowledge discovery, such as in hypothesis testing and variant prioritization in genetics studies. Moving forward, we hope frameworks like AMBER can be further developed to identify neural network architectures that are balanced in predictive power and interpretability.

## Methods

### Designing model search space

The AMBER neural architecture search framework consists of two components to design a child model for specific tasks: 1) a model search space with a large number of different child model architectures; and 2) a controller model that samples architectures from the model search space. For simplicity, we start by illustrating the design of model search space.

The model search space is a sequential collection of layers for the child model, where each layer has a number of candidate computational operations. More concretely, in this study, we aimed to design a 1D-convolutional neural network with 12 candidate convolutional layers. Each layer had 6 distinct computational operations: 1D convolution with filter size 4 or 8 (conv4, conv8), dilated 1D convolution with rate 10 and filter size 4 or 8 (dconv4, dconv8), max-pooling or average pooling with size 4 (maxpool, avgpool). These hyperparameters for computational operations were selected based on previous works^4,10^. Moreover, we added an identity mapping to each layer that maps input identically to output without any computations (identity), for potentially reducing the child model complexity. The twelve convolutional layers were connected to fixed input and output stem layers for inputs and outputs, respectively. We divided the 12 convolutional layers into 4 blocks of layers, where each block had doubled the number of filters from the previous block while reduced the size of the feature map by a factor of four. Layers within each block had identical number of filters. We set the first block to have 32 filters for searching architectures.

Formally, let the model space of *T*=12 layers be *Ω* = {*Ω*_1_,*Ω*_2_,…,*Ω*_*T*_} where *Ω*_*t*_ is the *t*-th layer. Under the current setup, *Ω*_*t*_ = {conv8, conv4, dconv8, dconv4, maxpool, avgpool, identity}, ∀ *t*. Let the selection of computational operations at *t*-th layer be a sparse categorical encoder, i.e. 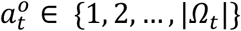. For example, 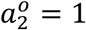 describes the operation for the second hidden layer of the child model is conv8. Therefore, child model computational operations are fully specified by a sequence of integers 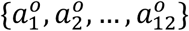; in total, different combinations of computational operations constitutes 8^12^ ≈ 6.9 × 10^10^ viable child models in the model space. The task of finding the child model computational operations can be subsequently considered as a multi-class classification problem with auto-regressive characteristics.

In addition to searching operations, we also incorporated the residual connections in the model search space. For the *t*-th layer, the residual connections from layers 1, 2, …, *t*-1 are binary encoded by 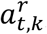,∀ *k* ∈ {1,2,..,*t* − 1}. If 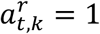, the residual connection is added from the output of the *k*-th layer to the *t*-th layer^13^. Having residual connections are essential for training deeper neural networks, but also significantly increases the complexity in architecture searching. For our 12-layer model space, residual connection search increased the search space by around 2^12×11/2^ ≈ 7.4 × 10^19^. Now with the residual connections, a full child model can be specified by a sequence of integers 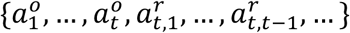 for brevity, we use *a*_*t*_ to denote both the operations and residual connections in the same layer and use {*a*_1_, …, *a*_*t*_} to represent the child model architecture.

### Efficient neural architecture search

We adopted Efficient Neural Architecture Search (ENAS) as the optimization method for searching the child network architectures in the model space^15^. ENAS employs a Recurrent Neural Network (RNN) as the controller model to sequentially predict the child model architecture from the model space. Briefly, the controller RNN, parameterized by *θ*, generates the child model architectures *a* with log-likelihood *π*(*a*; *θ*) and is trained by REINFORCE^37^. The policy gradient to maximize the reward *R_k_* over a batch of *m* sampled architectures is obtained by:

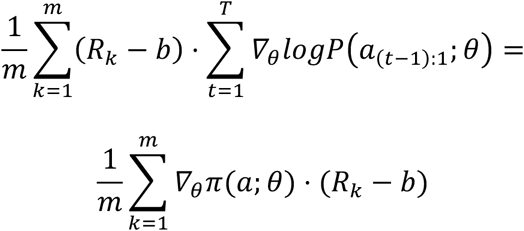

We set the reward *R*_*k*_ to be the validation AUROC of the *k*-th child model architecture; *b* is an exponential moving average of previous rewards to reduce the high variance of the policy gradient.

Another important feature that enables efficiently sampling of child architectures is the parameter sharing scheme among child models^15^. The computational graph for a child model is a Directed Acyclic Graph (DAG). Under the parameter sharing scheme, we build a large computational graph, named child DAG with parameters *ω*, which hosts all possible combinations of child model architectures. The key observation of ENAS is that each child model architecture is a subgraph of the child DAG, therefore the training of child model parameters is shared and significantly faster. The gradient for the child model parameters *ω* is obtained though Monte Carlo estimate:

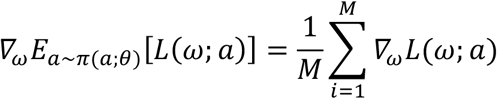

In this study, we made the following specifications and modifications in training the controller RNN parameters *θ* and the child DAG parameters *ω*. The controller RNN was parameterized as a 1-layer LSTM of 64 hidden units. Following the original ENAS implementation^15^, we set *M*=1 for updating *ω*; and regularized the proportion of residual connections if it deviated from 0.4. The child DAG was set according to the model space described in the previous section. The child DAG was first trained for a whole pass of the training data with a batch size of 1000 as a warm-up process. Next, the controller RNN sampled 100 child architectures from the child DAG and evaluated their rewards. The child architectures and the rewards were used to train the controller RNN parameters *θ*. Then we trained the child DAG with architectures sampled from updated *π*(*a*; *θ*). Both controller and child models were trained by Adam optimizer with a learning rate of 0.001. These two training processes were alternated for 300 iterations, and the child architecture with the best reward in the last controller step was extracted.

Sampled architectures were generated by sampling the computational operations uniformly and sampling the residual connections at the proportion of 0.4 as used in searched models. Finally, the child models of searched and sampled were trained from scratch to convergence using identical setup to facilitate downstream comparisons. Convergence was defined as validation AUROC not increasing for at least 10 epochs. To more robustly measure the accuracy of AMBER, we ran the search and sample processes for six times, respectively. Throughout the manuscript, all processing and analysis of searched and sampled models were strictly identical, except for how we derived their corresponding architectures. We referred to the searched model with best testing performance as AMBER-Seq and referred to the sampled model with best testing performance as AMBER-Base.

### Dataset for transcriptional regulatory activity prediction

The generic tasks of interest in this study were to predict transcriptional regulatory activity for a given DNA sequence. We aimed to design an end-to-end convolutional neural network model that takes raw one-hot encoded DNA as input. Following the previous work^4^, we used the pre-compiled training, validation and testing dataset downloaded from http://deepsea.princeton.edu/help/. The inputs were one-hot encoded matrices of DNA sequences built on hg19 reference human genome assembly. The training labels were compiled from a large compendium of publicly available ChIP-seq datasets, which measure the genome-wide molecular profiles such as protein binding or chemical modifications using high-throughput sequencing. In total, there are 919 distinct labels for ChIP-seq profiles of transcription factor binding, histone modification, and DNase accessibility assays in diverse human cell lines and tissues; and there are 4400000 training samples, 8000 validation samples and 455024 testing samples, each of 1000 bp (1000 x 4 when one-hot encoded) in length.

### Allele-specific binding analysis

A compendium of allele-specific transcription factor binding sites reported previously^24^ were compiled for benchmarking the variant effect predictions of the AMBER searched models. Briefly, ChIP-seq data were collected that measured genome-wide binding profiles for 83 unique transcription factors. For each binding site, binomial test was performed to test allelic imbalance and Benjamini-Hochberg False Discovery Rate (FDR) was used to correct for multiple testing. The baseline machine learning methods and the motif scorings were computed previously^24^. We further divided the variants into loss-of-binding alleles (reference reads ratio>0.6 and FDR<0.01), gain-of-binding alleles (reference reads ratio<0.4 and FDR<0.01), and neutral alleles (FDR>0.9).

The transcription factors were then mapped to the corresponding cell lines in the multi-tasking model. To benchmark the models of AMBER-Seq and AMBER-Base with other baseline models, we computed the variant effect scores as the log fold-change between reference allele prediction and alternative allele prediction, as previously described^4^. Then the AUROCs for distinguishing loss-of-function and gain-of-function alleles against the neural alleles were computed for each transcription factor from each model/motif, respectively. To compare the variant effect scores across different methods, we further rank-transformed the scores to the range of [-1, 1] while preserving scores at 0 for each method.

For the biological case study of variant effect prediction on SNP rs11658786, we reported its variant effect predictions from AMBER-Seq and AMBER-Base along with available baseline variant scoring methods^24^. Variants in high LD with the allele-specific variant of interest were queried from LDlink webserver^38^ (https://ldlink.nci.nih.gov/) using the EUR/CEU population and R^2^>0.9. Then the set of variants were processed by plink^39^ (v1.90) and plotted by R package gaston^40^. The eQTLs for allele-specific variants were queried using the GTEx web portal^41^ (https://www.gtexportal.org/home/).

### GWAS analysis

To evaluate the informativeness of the variant annotations from different model architectures, we used stratified LD-score regression^34^ to assess the heritability enrichment for variant annotations. First, we downloaded the summary statistics files from UK Biobank for disease phenotypes reported previously^35^. Selene^17^ (v0.4.2) was employed to process the genome-wide variant effect predictions for SNPs from the 1000 Genome Project (European cohort) for each transcriptional regulatory feature in both AMBER-designed AMBER-Seq model and uniformly-sampled AMBER-Base model. Then the variant effect predictions were subsequently converted to LD scores and regressed on the *χ*^2^ statistics using ldsc v1.0.1 Python implementation (https://github.com/bulik/ldsc), conditioned on a set of 97 baseline LD annotations from baselineLD v2.2 (https://data.broadinstitute.org/alkesgroup/LDSCORE/). We restricted our analyses for phenotypes with the ratio statistics less than 10% to avoid potential model misspecifications^34^. The enrichment P-values were computed by ldsc and corrected for multiple testing by Benjamini-Hochberg FDR. Regulatory features whose variant annotations were significant (FDR<0.05) in either the searched AMBER-Seq or the sampled AMBER-Base models were analyzed for their overlapping statistics and enrichment fold-changes across models.

## Data Availability

All data used in this study are publicly available and the URLs are provided in the corresponding sections in Methods.

## Code Availability

The AMBER package is available at GitHub: https://github.com/zj-zhang/AMBER; the analysis presented in this study is available at https://github.com/zj-zhang/AMBER-Seq

