## Supplementary Information for "An automated framework for efficiently designing deep convolutional neural networks in genomics"

### Supplementary Figures and Tables for An automated framework for efficiently designing deep convolutional neural networks in genomics

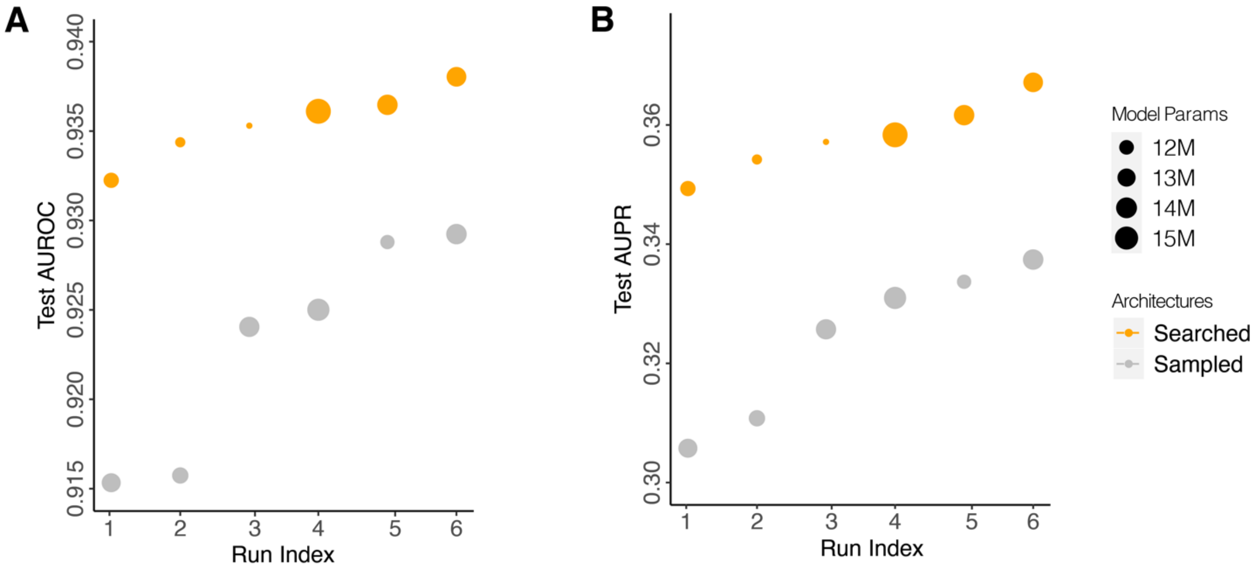


**Supplementary Figure 1**. An illustration of the total model parameters (dot sizes) for the transcriptional regulation prediction tasks with searched and sampled architectures. Models are ordered by their average prediction accuracy of all 919 tasks in **A**) AUROC and **B**) AUPR.

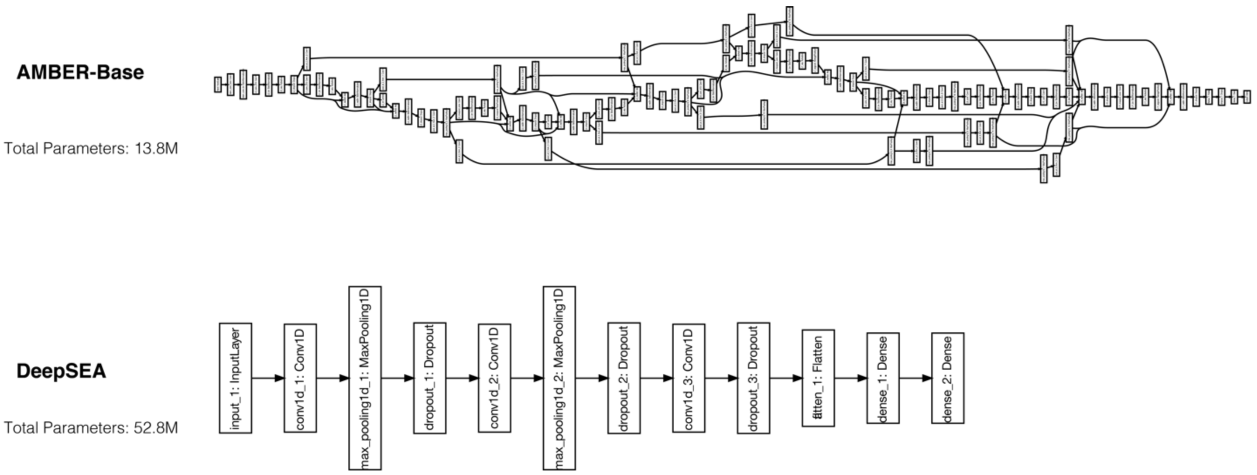


**Supplementary Figure 2**. Detailed model architectures for AMBER-Base (i.e. the best performing model with sampled architecture) and the original DeepSEA-2015 implementation, as compared to the AMBER-Seq architecture in Figure 2.

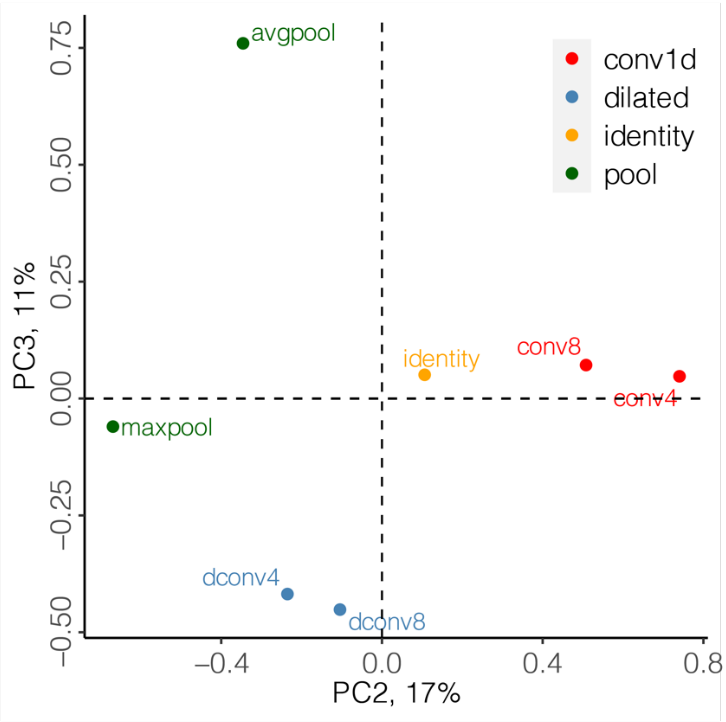


**Supplementary Figure 3**. Extended principal component analysis for the embedding vectors of different computational operations.

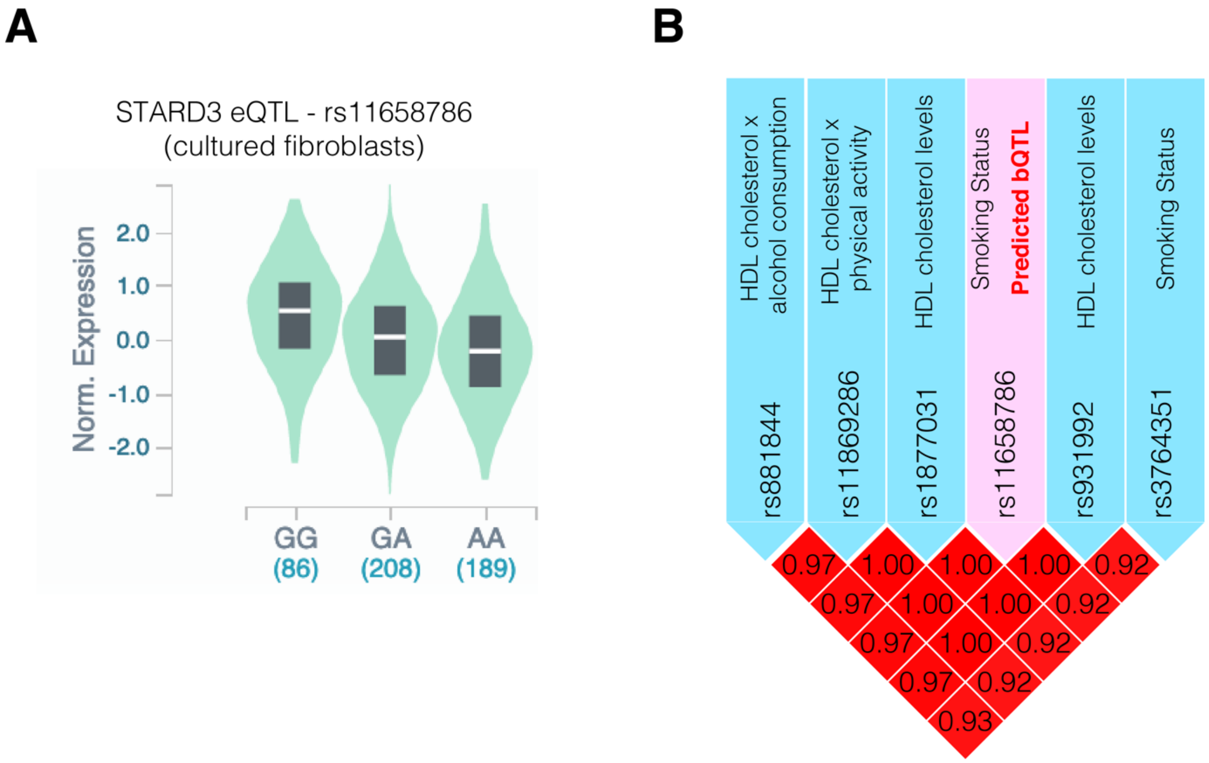


**Supplementary Figure 4**. **A)** eQTL and **B)** Linkage disequilibrium analysis for the biological case study SNP, rs11658786, as shown in Figure 4.

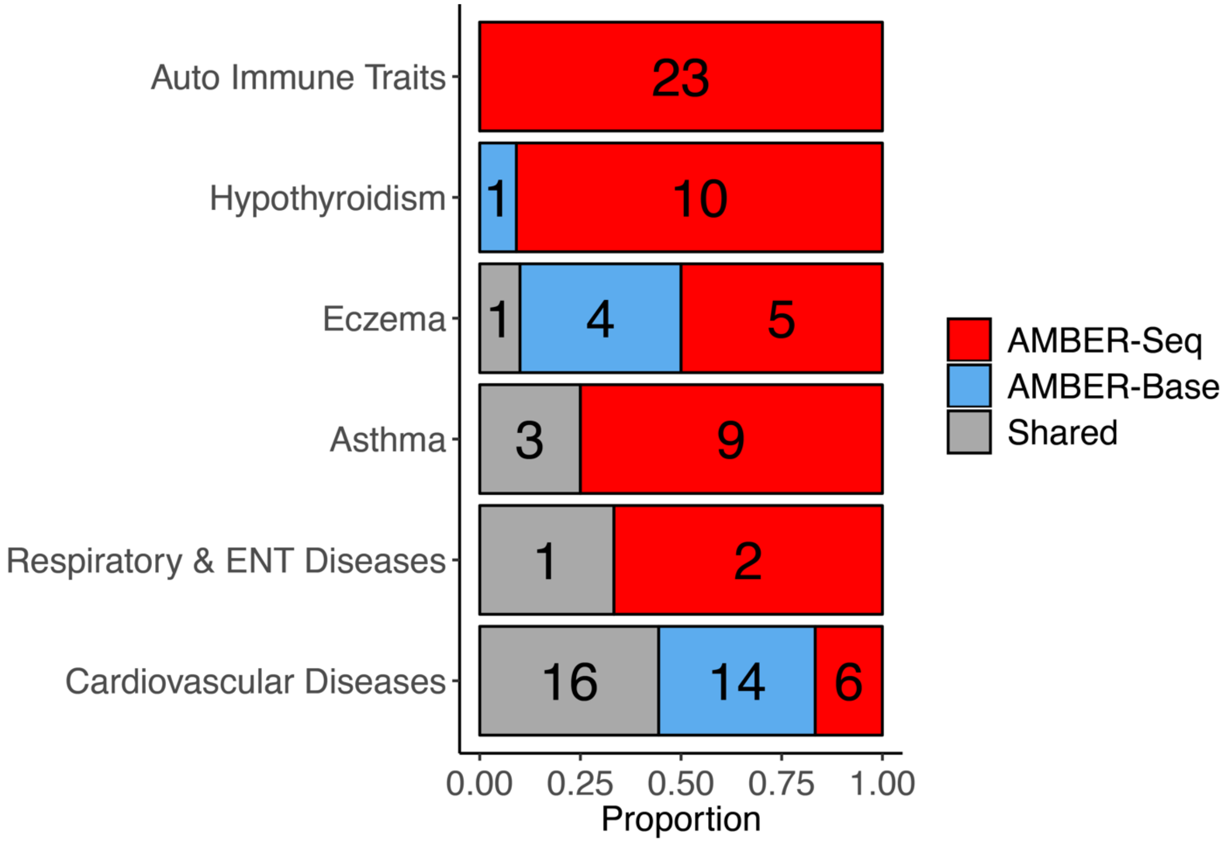


**Supplementary Figure 5**. The proportion and number of significant transcriptional regulatory features that were overlapped, unique for AMBER-Seq, or AMBER-Base models across GWAS disease phenotypes.

**Supplementary Table 1**. Detailed performance evaluations for AMBER searched, sampled models, and the original implementation by human experts.

| Run Index | Validation AUROC | Validation AUPR | Test AUROC | Test AUPR | Architecture |
|---|---|---|---|---|---|
| 6 | 0.945 | 0.420 | 0.938 | 0.367 | Search |
| 6 | 0.938 | 0.388 | 0.929 | 0.337 | Sample |
| 5 | 0.944 | 0.418 | 0.936 | 0.362 | Search |
| 5 | 0.938 | 0.389 | 0.929 | 0.334 | Sample |
| 4 | 0.934 | 0.382 | 0.925 | 0.331 | Sample |
| 4 | 0.943 | 0.400 | 0.936 | 0.358 | Search |
| 3 | 0.932 | 0.372 | 0.924 | 0.326 | Sample |
| 3 | 0.942 | 0.413 | 0.935 | 0.357 | Search |
| 2 | 0.925 | 0.360 | 0.916 | 0.311 | Sample |
| 2 | 0.942 | 0.409 | 0.934 | 0.354 | Search |
| 1 | 0.926 | 0.355 | 0.915 | 0.306 | Sample |
| 1 | 0.942 | 0.405 | 0.932 | 0.349 | Search |
| 0 | 0.937 | 0.402 | 0.931 | 0.338 | Human |
